# Novel pectin from crude polysaccharide of *Syzygium aromaticum* against SARS-CoV-2 activities by targeting 3CLpro

**DOI:** 10.1101/2021.10.27.466067

**Authors:** Can Jin, Bo Feng, Rongjuan Pei, Yaqi Ding, Meixia Li, Xia Chen, Zhenyun Du, Yangxiao Ding, Chunfan Huang, Bo Zhang, Xinwen Chen, Yi Zang, Jia Li, Kan Ding

**Affiliations:** College of Pharmacy, Nanjing University of Chinese Medicine, 138 Xianlin Avenue, Nanjing, Jiangsu Province 210029, China; Glycochemistry and Glycobiology Lab, Key Laboratory of Receptor Research; National Center for Drug Screening, State Key Laboratory of Drug Research, Shanghai Institute of Materia Medica, Chinese Academy of Sciences, 555 Zu Chong Zhi Road, Shanghai 201203, P. R. of China; University of Chinese Academy of Science, No.19A Yuquan Road, Beijing 100049, P. R. China; State Key Laboratory of Virology, Wuhan Institute of Virology, Center for Biosafety Mega-Science, Chinese Academy of Sciences, Wuhan, 430071, China; Zhongshan Institute for Drug Discovery, Shanghai Institute of Materia Medica, Chinese Academy of Science. SSIP Healthcare and Medicine Demonstration Zone, Zhongshan Tsuihang New District, Zhongshan, Guangdong, China, 528400; Henan Polysaccharide Research Center, Academy of Chinese Medical Sciences, Henan University of Chinese Medicine, Zhengzhou 450046, Henan, China; Shanghai High School International Division, No.989, Baise Road, Shanghai, China, 200231

**Keywords:** COVID-19, SARS-CoV-2, 3CL protease, PLpro, RdRp, Syzygium aromaticum, Polysaccharide, Pectin

## Abstract

To date, COVID-19 is still a severe threat to public health, hence specific effective therapeutic drugs development against SARS-CoV-2 is urgent needed. 3CLpro and PLpro and RdRp are the enzymes required for the SARS-CoV-2 RNA synthesis. Therefore, binding to the enzyme may interfere the enzyme function. Before, we found that sulfated polysaccharide binding to 3CLpro might block the virus replication. Hence, we hypothesize that negative charged pectin glycan may also impede the virus replication. Here we show that 922 crude polysaccharide from *Syzygium aromaticum* may near completely block SARS-CoV-2 replication. The inhibition rate was 99.9% (EC_50_ : 0.90 μM). Interestingly, 922 can associates with 3CLpro, PLpro and RdRp. We further show that the homogeneous glycan 922211 from 922 may specifically attenuate 3CL protease activity. The IC_50_s of 922 and 922211 against 3CLpro are 4.73 ± 1.05 µM and 0.18 ± 0.01 µM, respectively. Monosaccharide composition analysis reveals that 922211 with molecular weight of 78.7 kDa is composed of rhamnose, galacturonic acid, galactose and arabinose in the molar ratio of 8.21 : 37.81 : 3.58 : 4.49. The structure characterization demonstrated that 922211 is a homogalacturonan linked to RG-I pectin polysaccharide. The linear homogalacturonan part in the backbone may be partly methyl esterified while RG-I type part bearing 1, 4-linked α-Gal*p*A, 1, 4-linked α-Gal*p*AOMe and 1, 2, 4-linked α-Rha*p*. There are four branches attached to C-1 or C4 position of Rhamnose glycosyl residues on the backbone. The branches are composed of 1, 3-linked β-Gal*p*, terminal (T)-linked β-Gal*p*, 1, 5-linked α-Ara*f*, T-linked α-Ara*f*, 4-linked α-Gal*p*A and/or 4-linked β-Gal*p*A. The above results suggest that 922 and 922211 might be a potential novel leading compound for anti-SARS-CoV-2 new drug development.

## 1. Introduction

The severe acute respiratory syndrome coronavirus 2 (SARS-CoV-2) is kind of novel coronavirus. It made a serious life-threatening disease Coronavirus Disease 2019 (COVID-19). The first case of COVID-19 was reported by the Work Health Organization (WHO) on December 31, 2019 (Majumder & Minko, 2021). Owing to SARS-CoV-2 infection spreading rapidly and a shortage of specific treatments for COVID-19, it has made an ongoing pandemic in a growing number of countries, which has caused a serious public health threat (Laventhal et al., 2020). On March 11, 2020, WHO declared COVID-19 as a global pandemic (Cucinotta & Vanelli, 2020). Up to now, the virus pandemic in more than 200 countries has resulted in major loss of human life globally. As of October 19, 2021, more than 240 million people were infected with the COVID-19 disease and 4.9 million deaths have been reported globally and the number is increasing rapidly (**https://covid19.who.int/**). Since the outbreak of COVID-19, a mass of studies have attempted to reveal the structural characteristics of SARS-CoV-2 and its infection mechanism. Some investigations found that SARS-CoV-2 is an enveloped virus consisting of a positive-sense, single-stranded RNA genome of around 30 kb (Asselah et al., 2021). The envelope is covered with glycoprotein spikes. SARS-CoV-2 viral genome encoded its structural proteins namely, spike (S) surface glycoprotein, membrane (M) protein, envelope (E) protein, and nucleocapsid (N) protein and nonstructural proteins (RNA polymerase, RdRp; papain-like protease, PLpro; coronavirus main protease, 3CLpro) (Yadav et al., 2021). SARS-CoV-2 and SARS-CoV are highly similar genetically and at the protein production level (Hatmal et al., 2020). Almost 85% homology of this virus is similar to the SARS-CoV (Petrosillo et al., 2020). Similar to SARS-CoV, the S1 protein on the surface of the SARS-CoV-2 binds to ACE2 on the plasma membrane of infected cells, initiating receptor-mediated endocytosis (Yeung et al., 2021). Similarly, in the SARS-CoV-2 replication cycle, viral proteinases 3CLpro and PLpro cleave viral polyproteins into effector proteins to ensure normal replication (Moustaqil et al., 2021). As early as the study of coronavirus, scientists found that 3CL and PLpro proteins were attractive target molecules for the treatment of coronavirus, because they are key enzymes in the process of virus replication (Báez-Santos et al., 2015). These mechanisms suggest that the development of anti-COVID-19 drugs and vaccines induced antibody can inhibit the binding of S1 protein to host cell ACE2 and target 3CL and PLpro proteins (Arvin et al., 2020; Yan & Gao, 2021). While the best interventions to control and ultimately stop the pandemic are prophylactic vaccines, antiviral therapeutics are important to limit morbidity and mortality in those already infected (Froggatt et al., 2020). Through structure-assisted drug design, virtual drug screening and high-throughput screening, scientists found that natural and synthetic active substances had the potential to become the lead compounds for the new drug development for the treatment of COVID-19. It is noteworthy that some natural small molecules, such as flavonoids can target 3CL and PLpro proteins (Russo et al., 2020) and inhibit the binding of S1 and ACE2 (Mouffouk et al., 2021). They inhibit the replication of SARS-CoV-2 in Vero E6 cells (Muchtaridi et al., 2020). Because of its non-toxic and multi-target characteristics, macromolecules polysaccharides have attracted the attention of scientists in the treatment of various diseases (Kocabiyik et al., 2021). In fact, scientists have also explored that polysaccharides have good anti-SARS-CoV-2 effects, such as heparin, some seaweed polysaccharides, and sulfated derivatives of chitosan (Chen et al., 2020; Modak et al., 2021; Pereira & Critchley, 2020; Tandon et al., 2021). The similarity of these polysaccharides is that they contain a large amount of sulfate ions. Likewise, it was reported that Heparin blocked SARS-CoV-2 binding and infection in mechanism of negatively charged sulfate and carboxyl groups in Heparin stabilizing the association with several positively charged amino acid residues of Spike (Hu et al., 2021). Hence, we hypothesize that natural pectin polysaccharides also with negative charged with a mass of carboxyl groups might also effectively inhibit SARS-CoV-2. To address, flowers of the traditional Chinese medicine *Syzygium aromaticum* L. were selected firstly as the source of pectin polysaccharides extraction in this study. *Syzygium aromaticum* belongs to the genus Eupatorium. Actually, flower buds and fruits of *Syzygium aromaticum* is often employed as medicine for diseases treatment in China and Southeast Asia. Ancient medical books and modern pharmacological studies have shown that *Syzygium aromaticum* has strong catharsis, insecticidal and bacteriostatic effects (Batiha et al., 2019, 2020; Radünz et al., 2019). However, studies have shown that these biological activities often arise from small molecules in *Syzygium aromaticum*, such as eugenol, volatile oil, etc. In this paper, we firstly extracted and isolated polysaccharide 922 from *Syzygium aromaticum*. Then we test the bioactivity of the polysaccharide against SARS-CoV-2 activity. Further, one homogeneous polysaccharide 922211 was purified from 922 followed by 3CPpro, PLpro and RdRp enzymes activities measurement using this polysaccharide and its native one. Then we characterized the structure of one homogeneous polysaccharide 922211 using the method combining chemical and spectral analysis, including methylation analysis, partial acid hydrolysis, GC-MS (Gas chromatography-mass spectrometry) and nuclear magnetic resonance (NMR) spectroscopy.

## 2. Experimental

### 2.1. Materials and reagents

Dried flower buds of *Syzygium aromaticum*. were purchased from Bozhou Decoction Pieces and Medicinal Materials Factory (Bozhou, Anhui Province, China). DEAE Sepharose Fast Flow and Sephacryl S-300 HR was obtained from GE healthcare (Danderyd, Sweden, USA). CMC (1-Cyclo-hexyl-3- (2-morpholinoethyl) carbodiimide metho-p-toluenesulfonate) and Iodomethane were purchased from TCI (Tokyo, Japan). T-series Dextrans were obtained from Amersham Pharmacia Biotech (Little Chalfont, Buckinghamshire, UK). Standard monosaccharides were bought from Shanghai Macleans Biochemical Technology Co., Ltd. (Shanghai, China). BCA kit was obtained from Shanghai biyuntian Biotechnology Co., Ltd (Shanghai, China). Other reagents were analytical grade and from Sinopharm Chemical Reagent Co. Ltd. (Shanghai, China).

### 2.2. Extraction, isolation and purification of polysaccharides

The crude polysaccharide was extracted through water extraction and alcohol precipitation. In brief, the dried flower buds of *Syzygium aromaticum* was immersed (solid-liquid ratio : 1 kg/15 L) in water for 12 h. The soaked mixture was extracted with boiling water for 2 h, twice in total. The combined supernatant was concentrated, dialyzed (dialysis membrane for small molecule), concentrated, centrifuged and precipitated with three volumes of 95% EtOH. The crude polysaccharide 922 (8 g) was dissolved in 100 mL distilled water and centrifuged (4000 g, 10 min/time). The supernatant was fractionated by anion-exchange chromatography on DEAE-cellulose column (Cl^-^, 50 cm × 5 cm), eluted stepwise with distilled water, 0.05, 0.1, 0.2 and 0.4 M NaCl solution. For the sample solution collected by each mobile phase, two curves need to be drawn. One is the polysaccharide elution curve detected by phenol-sulfuric acid method and the other is the protein curve determined by BCA kit. Hence, based on the curves, the corresponding polysaccharide and protein from one elution would be accumulated. The Elution processes was shown in **Fig. 6F**. Among them, the fraction eluted with 0.2 M NaCl elution was collected, concentrated and lyophilized to obtain polysaccharide 9222. Subsequently, 9222 (200 mg) was dissolved in 5 mL distilled water and centrifuged (4000 g, 10 min/time). The supernatant was further purified by gel permeation chromatography using Sephacryl S-300 column (100 cm × 2.6 cm) and the Sephacryl S-100 column (100 cm × 2.6 cm), by which was eluted with 0.2 M NaCl to achieve the target polysaccharide 922211. The relative molecular weight of 922211 was estimated by high performance gel permeation chromatography (HPGPC) with series-connected Shodex SUGAR KS-804 and Shodex SUGAR KS-802 columns.

### 2.3. Homogeneity and molecular weight determination

Determining the homogeneity and molecular weight of polysaccharides were measured by high performance gel permeation chromatography (HPGPC) on an Agilent 1260 HPLC system equipped with series-connected Shodex SUGAR KS-804 and Shodex SUGAR KS-802 columns, with 0.1 M NaNO_3_ used as the mobile phase at a flow rate of 0.5 mL/min (Cong et al., 2014). All samples were prepared as 4 mg/mL in mobile phase, and 10 μL of solution was injected in each run (60 min for each run). The eluate was monitored with an RI (Keep in 25 °C) and a UV detector, and the column temperature was kept at 35 °C.

### 2.4. Monosaccharide composition analysis

The monosaccharide composition was analyzed using PMP pre-column derivatization based on the previous reported (J. Dai et al., 2010). In briefly, 922211 (2 mg) was hydrolyzed with 4 mL of 2 M TFA (trifluoroacetic acid), followed by PMP derivation. 10 μL of the derivative solution was analyzed by high performance liquid chromatography (HPLC) to understand the sugar composition.

### 2.5. NMR analysis

For NMR analysis, 922211 (30 mg) was deuterium-exchanged and dissolved by 0.5 mL D_2_O (99.8% D), and then lyophilized and redissolved in 0.5 mL D_2_O (99.8% D). The ^1^H, ^13^C NMR and 2D NMR spectra (COSY, HSQC and HMBC) were measured at 25 °C with acetone as internal standard (δH = 2.29, δC = 31.5). NMR spectra were recorded on a Bruker AVANCE III NMR spectrometer.

### 2.6. Methylation analysis

The dried polysaccharide (10 mg) was methylated for 3-4 times based on previous methods (HAKOMORI, 1964). The methylated polysaccharide was hydrolyzed and then reduced with sodium borohydride and acetylated. The partially methylated alditol acetates were examined by gas chromatography–mass spectrometry (GC-MS). Mass spectra of the derivatives were analyzed using Complex Carbohydrate Structural Database of Complex Carbohydrate Research Centre (http://www.ccrc.uga.edu/).

### 2.7. Uronic acid reduction

The approach of uronic acid reduction was based on the reported method (Taylor & Conrad, 1972). In brief, 40 mg polysaccharide was dissolved in 40 mL H_2_O. CMC (600 mg) was added and pH was kept at 4.75 with 0.01 M HCl for 2 h. Then 2 M fresh aqueous sodium borohydride (16 mL) was added slowly to the mixture (Sodium borohydride solution shall be added within 30-60 min) and maintained pH at 7 with 4 M HCl for 2 h at room temperature. The mixture was dialyzed (1,000 mL × 4) for 24 h at room temperature. Then the retentate was lyophilized to achieve carboxyl reduced polysaccharide, followed by monosaccharide composition and glycosyl residues analyses.

### 2.8. Enzymatic activity and inhibition assays

The enzyme activity and inhibition assays of SARS-CoV-2 3CLpro have been described previously (W. Dai et al., 2020; Jin et al., 2020). Briefly, the recombinant SARS-CoV-2 3CLpro (40 nM at a final concentration) was mixed with each compound in 50 μL assay buffer (20 mM Tris, pH7.3, 150 mM NaCl, 1mM EDTA, 1% Glycerol, 0.01% Tween-20) and incubated for 10 min. The reaction was initiated by adding the fluorogenic substrate MCA-AVLQSGFRK (DNP) K (GL Biochem, Shanghai), with a final concentration of 20 μM. After that, the fluorescence signal at 320 nm (excitation)/405 nm (emission) was immediately measured by continuous 8 points for 8 min with an EnVision multimode plate reader (Perkin Elmer, USA). The initial velocity was measured when the protease reaction was proceeding in a linear fashion.

The activity of SARS-CoV-2 PL^pro^ was also measured by a continuous 8 points fluorometric assay for 8 min. Briefly, the recombinant SARS-CoV-2 PL^pro^ (40 nM at a final concentration) was mixed with each compound in 50 μL assay buffer (20 mM Tris pH 8.0, 0.01% Tween 20, 0.5 mM DTT) and incubated for 10 min. The reaction was initiated by adding the substrate Z-RLRGG-AMC (GL Biochem, Shanghai) with a final concentration of 50 μM, using wavelengths of 355 nm and 460 nm for excitation and emission, measured by an EnVision multimode plate reader (Perkin Elmer, USA).

The detection of RNA synthesis by SARS-CoV-2 RdRp complex were established based on a real-time assay with the QuantiFluor® dsDNA Dye (Promega), which contains a fluorescent DNA-binding dye that enables sensitive quantitation of small amounts of double-stranded DNA (dsDNA) in solution. The fluorescence was measured using wavelengths of 504 nm and 531 nm for excitation and emission, measured by an EnVision multimode plate reader (Perkin Elmer, USA). The assay records the synthesis of dsRNA in a reaction using a poly-U molecule as a template and ATP as the nucleotide substrate. Reactions were performed in individual wells of white 384-well low volume round bottom plates. The standard reaction contained 50 mM Tris-HCl, pH 7.5, 50 mM Ammonium acetate, 0.5 mM MnCl_2_, 20 μM ATP, 0.2 μM poly-U template–primer RNA, 0.01% Tween-20

### 2.9. ELISA

10 µg/mL ACE2 coating buffer were used to treat the 96 well plate at 4 °C overnight following with 200 µL washing buffer for three times. Then the 96-well plate was blocked by 2% BSA at room temperature for 2 h. After that, 100 µL biotinylated S1 protein was added and incubated at room temperature. At the same time, the positive control and the negative control were set. After incubation for 1 h, the plate was washed for three times and each time for 5 min. Subsequently, 100 µL Streptavidin-HRP was added to final concentration of 200 ng/mL at room temperature and incubated for 1 h. After incubation for 1 h, the plate was washed for three times. Then, 100 µL TMB were added and incubated in the dark for 35 min. Finally, 50 µL stop solution were added to stop the reaction followed by detection at 450 nm by microplate reader (BioTek).

### 2.10. Antiviral test in vitro

The experiments related to SARS-CoV-2 are completed at National Biosafety Laboratory, Wuhan, Chinese Academy of Sciences. SARS-CoV-2 (WIV04) was passaged in Vero E6 cells and tittered by plaque assay. Vero E6 cells were treated with polysaccharides or positive control at indicated concentration and infected by SARS-CoV-2 virus at MOI 0.01. After 24 h incubation at 37 °C, supernatants were collected and the viral RNAs were extracted by Magnetic Beads Virus RNA Extraction Kit (Shanghai Finegene Biotech, FG438), and quantified by real-time RT-PCR with Taqman probe targeting to the RBD region of S gene.

## 3. Results and discussion

### 3.1. Purity, Molecular weight and monosaccharide composition analysis of 922211

The crude polysaccharide 922 was achieved at a yield of 4.2% (42 g/kg) by water extraction. 922 was further fractioned by DEAE Sepharose™ Fast Flow to obtain 9222 component (386.4 mg/8 g, yield is 4.83%) from the 0.2 M NaCl eluent. 9222 was further purified through gel permeation chromatography to achieve polysaccharide 922211 (61.28mg/200 mg, yield is 30.64%). Polysaccharide 922211 homogeneity was estimated by HPGPC, in which it showed one symmetrical peak. The relative molecular weight of 922211 was estimated to be 78.7 kDa. Monosaccharide composition analysis showed that 922211 was composed of rhamnose, galacturonic acid, galactose and arabinose in the molar ratio of 8.21 : 37.81 : 3.58 : 4.49 (**Table. 1**).

**Table. 1.** Monosaccharide composition of 922211, its hydrolysate, and reduced derivatives.

| Monosaccharides | Molar ratios (%) |  |  |  |
| --- | --- | --- | --- | --- |
|  | 922211 | R3922211 | P2922211I | R3P2922211I |
| Rha | 8.21 | 15.05 | 5.30 | 1.96 |
| GalA | 37.81 | nd | 47.07 | nd |
| Gal | 3.58 | 64.91 | trace | 49.89 |
| Ara | 4.49 | 8.87 | nd | nd |

nd: not detected.

### 3.2 Linkage pattern analysis

To determine the glycosyl linkage type, 922211 was methylated, hydrolyzed, reduced and acetylated to produce the partially methylated alditol acetates (PMAA), which was analyzed by gas chromatography-mass spectrometry (GC-MS) (**Table 2**). For 922211, the linkage pattern of the galactose residues included 1, 3-linked Gal (7.81%) and Terminal (T)-linked Gal (12.67%). The arabinose residues in the 922211 were consisted of Terminal (T)-linked Ara (16.46%) and 1, 5-linked Ara (4.80%). The rhamnose residues contained 1, 2, 4-linked Rha (19.07%). 922211 had a mass of galacturonic acid (**Table 1**), which could not be methylated successfully. Hence, the carboxyl group of galacturonic acid needs to be reduced to hydroxyl to acetylate and produce the PMAA successfully to then analyze the linkage type of galacturonic acid in 922211. Noticeably, a new linkage style 1, 4-linked Gal (37.91%) appeared in reduced polysaccharide R3922211 after being methylated. Combined with the monosaccharide composition analysis of 922211 (galacturonic acid 37.81%), it could be inferred that the galacturonic acid residues of 922211 is 1, 4-linked GalA.

**Table. 2.**
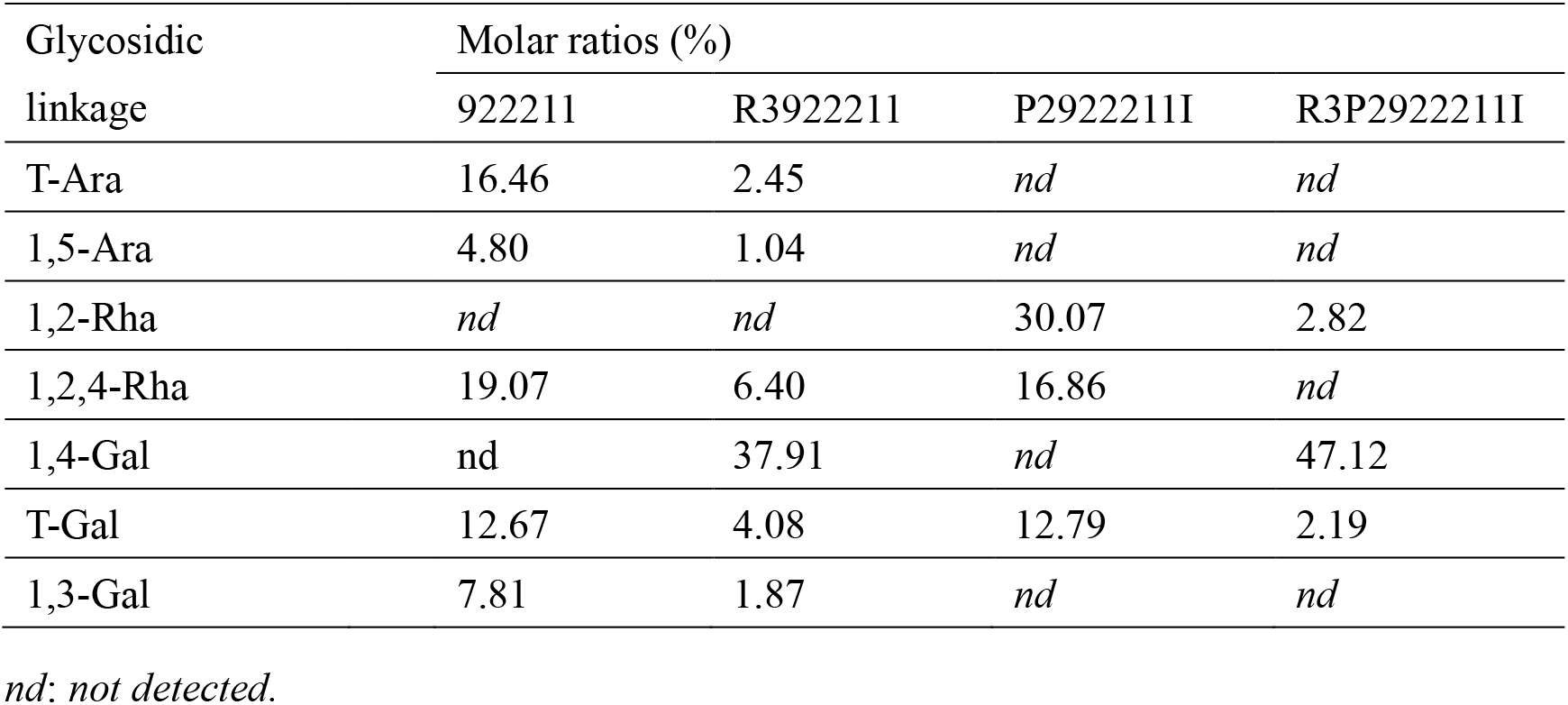
Linkage styles of 922211 and its hydrolyzed and reduced derivatives.

### 3.3 Partial acid hydrolysis

To elucidate the detail structure features of the backbone and branches of the polysaccharide, 922211 was subjected firstly to partial acid hydrolysis, followed by dialysis against de-ionized water. The intra-dialysate P2922211I (108 mg, yield: 72%) was obtained. Then the homogeneity of P2922211I was analyzed by HPGPC and identified to be a homogenous fraction with an average molecular weight of 41.4 kDa. Monosaccharide composition analysis revealed that P2922211I was composed of Rha and GalA in approximately molar ratio of 5.30 : 47.07. Comparing with 922211, the relative amount of galacturonic acid in P2922211I increased while the relative amount of galactose and arabinose disappeared, indicating that galactose and arabinose might locate in the side chains. Methylation analysis results suggested that P2922211I was composed of T-linked Gal (12.79%), 1, 2-linked Rha (30.07%) and 1, 2, 4-linked Rha (16.86%). Since P2922211I was acid polysaccharide, to make sure where the carboxyl group came from why residue, this polysaccharide was reduced to obtain R3P2922211I. Comparing the amount of sugar residues of P2922211I and R3P2922211I, we found that a new linkage style 1, 4-linked Gal (47.12%) appeared. This result helped to deduce that the galacturonic acid residue of P2922211I was 1, 4-linked GalA.

Since sugar composition analysis indicated that P2922211I contained Rha and GalA in the ratio of 5.30 : 47.07, while the methylation results show the residues linkage type of this fraction has 1,2-linked (30.07%), l,2,4-linked (16.86) and T-Gal (12.79) without any GalA linkage type due to the methylation method defect. However, after the reduction, the methylation results demonstrated that R3P2922211I had 1,4-linked Gal (47.12%) represented GalA as dominant part in the P2922211I and trace 1,2-linked Rha (2.82) and T-Gal (2.19). The above results suggested that backbone of 922211 somehow might contain homogalacturonan region of pectin at least in P2922211I fraction. The branched chains were composed of T-, 1, 3-linked Gal and T-, 1, 5-linked Ara. This inference was further evidenced by 1D and 2D NMR data analysis as followed.

### 3.4. NMR spectral analysis

The ^13^C (**Fig. 1A, C**) and ^1^H (**Fig. 1B**) NMR spectra were assigned and depicted in **Table 3** supported by the monosaccharide composition, methylation analysis, partial acid hydrolysis, two-dimension NMR spectra of 922211 and P2922211I (**Fig. 2**).

**Table. 3.**
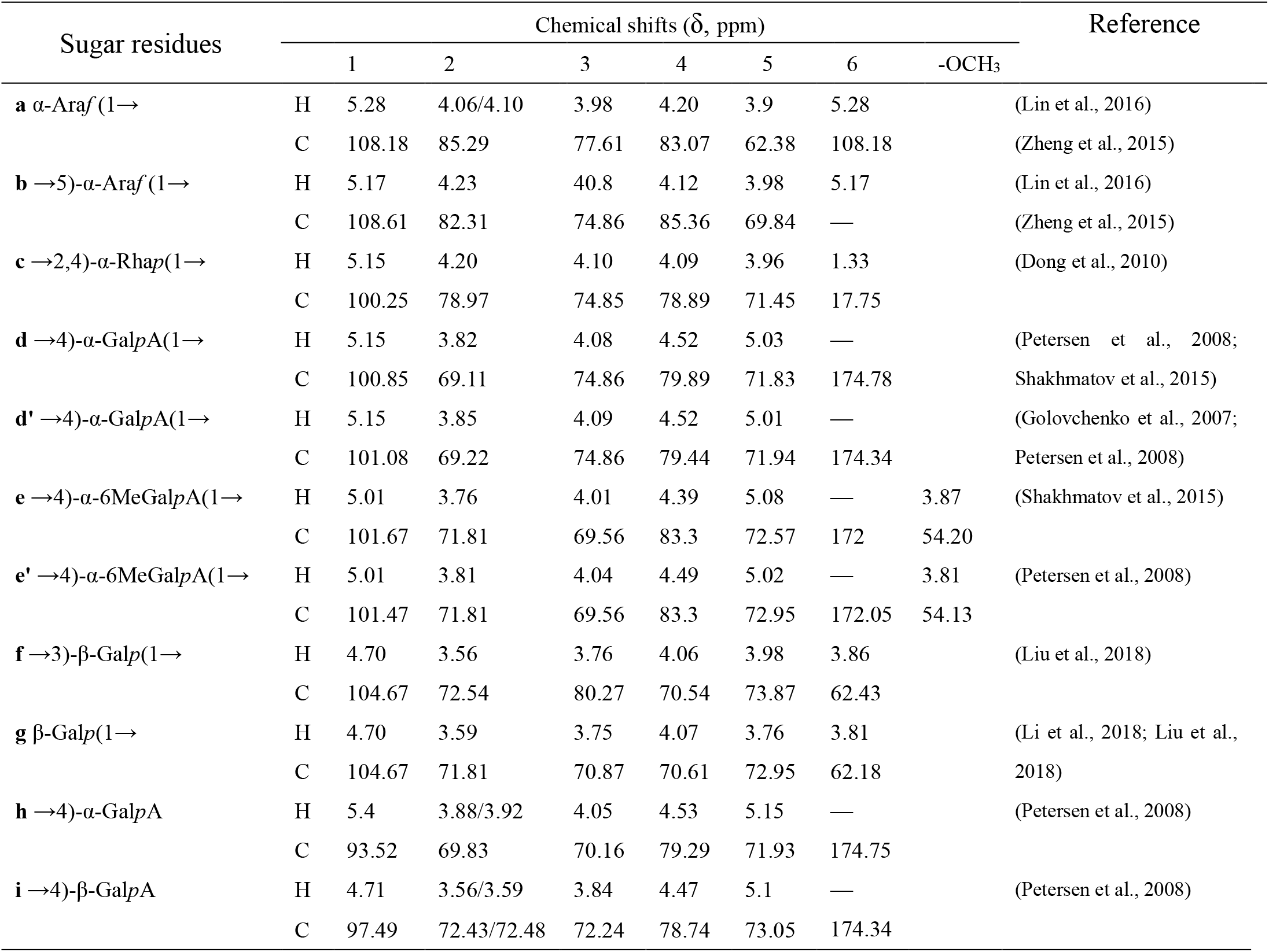
^1^H and ^13^C NMR chemical shifts (ppm) assignments for major signals of 922211.

**Fig. 1.**
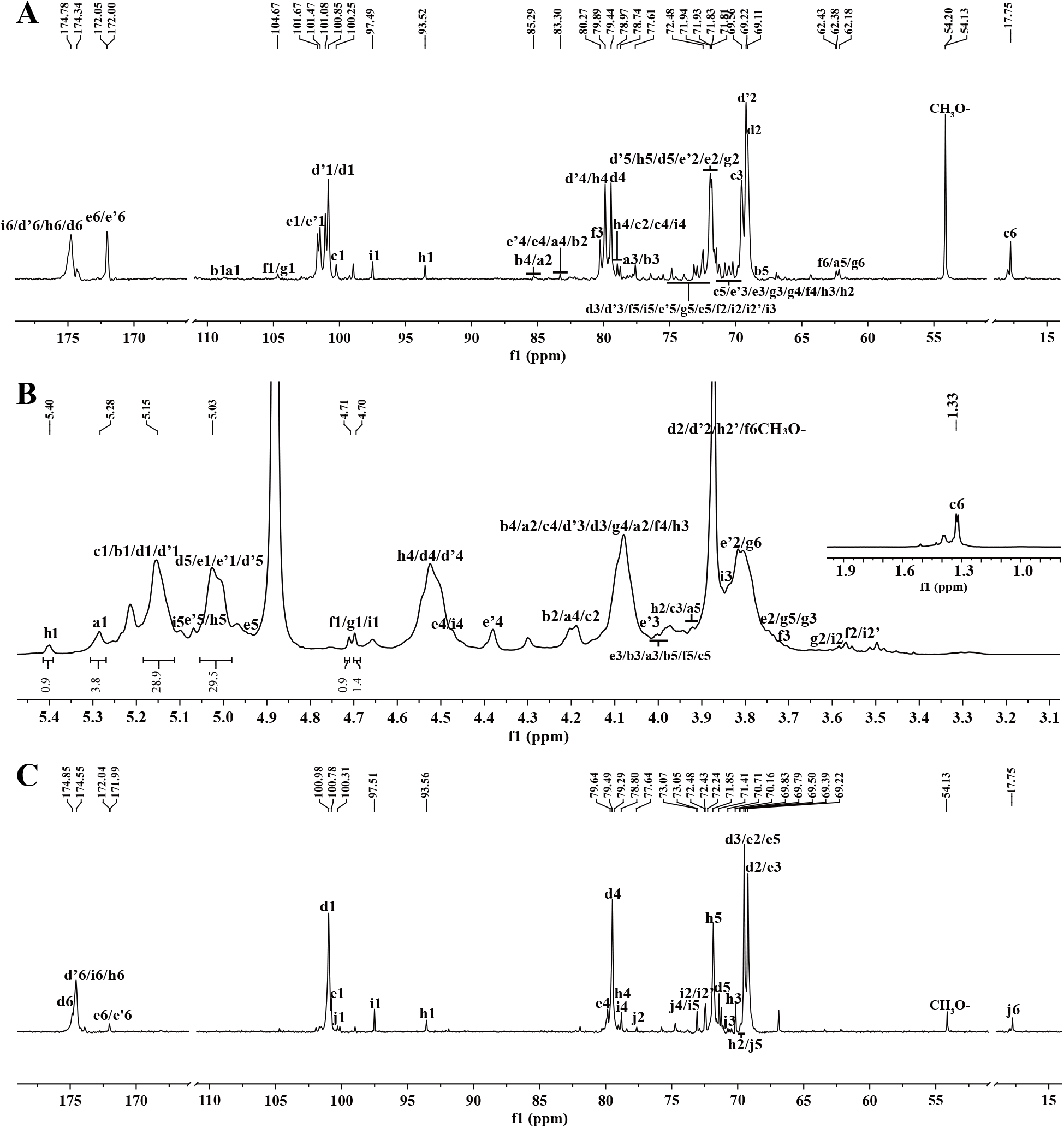
^13^C NMR spectra of the polysaccharide 922211 and its degraded polysaccharide P2922211I A: ^13^C NMR spectrum of 922211; B: ^1^H NMR spectrum of 922211; C: ^13^C NMR spectrum of P2922211I (**a**. T-linked a-Ara*f*; **b**. 1,5 -linked a-Ara*f*; **c**. 1, 2, 4-linked a-Rha*p*; **d/d’**. 1, 4-linked a-Gal*p*A; **e/e’**. 1, 4-linked α-6MeOGal*p*A; **f**. 1, 3-linked β-Gal*p*; **g**. T-linked β-Gal*p*; **h**. 4-linked a-Gal*p*A; **i**. 4-linked β-Gal*p*A; **j**. 1, 2-linked a-Rha*p*).

**Fig. 2.**
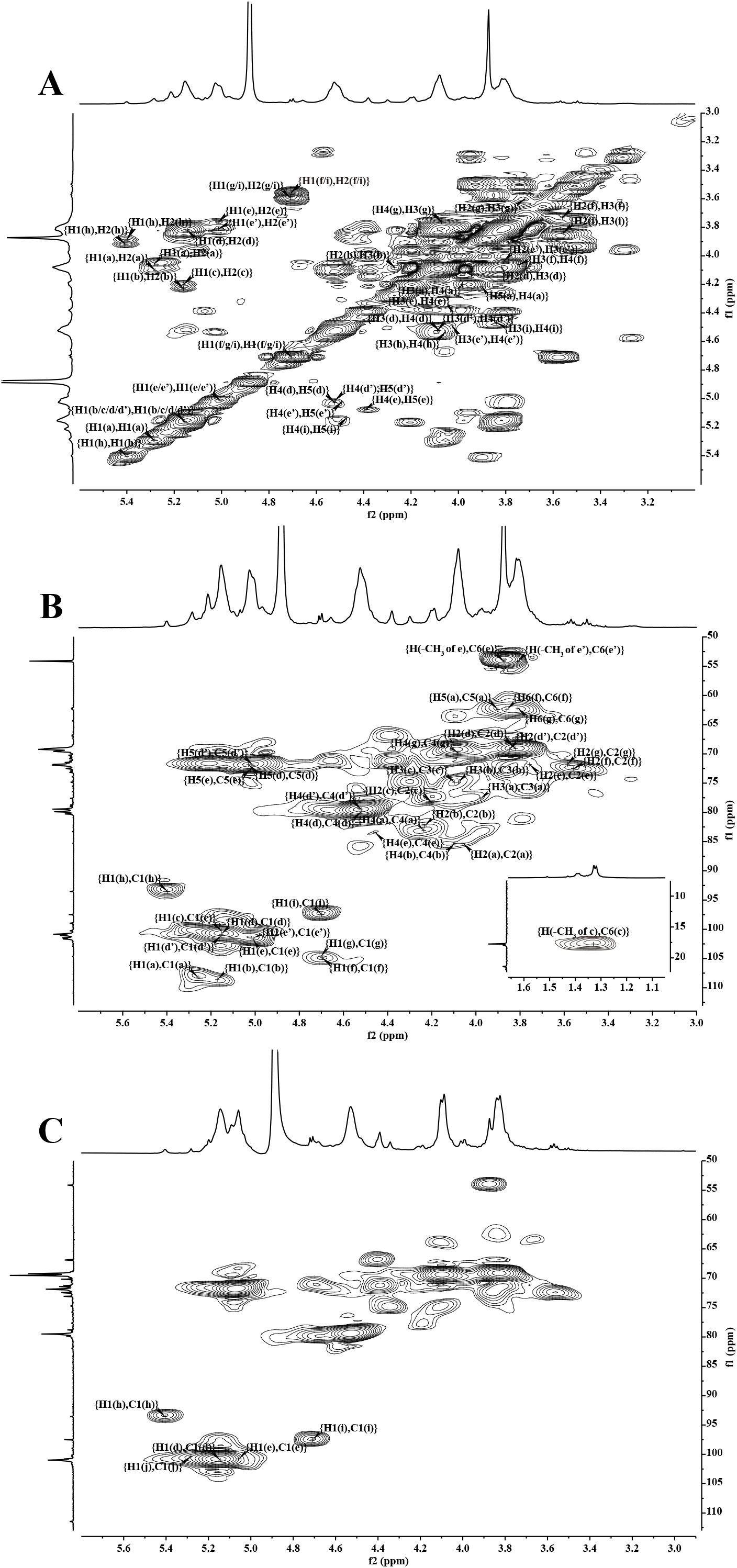
COSY (A) and HSQC (B) spectra of 922211 (**a**. T-linked a-Ara*f*; **b**. 1,5 -linked a-Ara*f*; **c**. 1, 2, 4-linked a-Rha*p*; **d/d’**. 1, 4-linked a-Gal*p*A; **e/e’**. 1, 4-linked α-6MeOGal*p*A; **f**. 1, 3-linked β-Gal*p*; **g**. T-linked β-Gal*p*; **h**. 4-linked a-Gal*p*A; **i**. 4-linked β-Gal*p*A; **j**. 1, 2-linked a-Rha*p*).

In ^13^C NMR spectrum, the two intense anomeric signals at δ 100.85 and δ 101.08 could be assigned to C-1 of 1, 4-linked α-Gal*p*A **(d, d’)** in different chemical environments, respectively. Signals at δ 101.47 and δ 101.67 could be assigned to C-1 of 1, 4-linked α-6MeOGal*p*A **(e’, e)** in different chemical environments, respectively. Indeed, the existence of methoxy group was further proved by the appearing signals at δ 172.00 and δ 172.05, which prompting partial Gal*p*A residues might exist methyl ester. Furthermore, signals at δ 54.13 and δ 54.20 indicated that methoxy group was attached to the C-6 position of 1, 4-linked α-6MeOGal*p*A. Signals at δ 93.52 and δ 97.49 might be attributed to anomeric carbon of 4-linked α-Gal*p*A and 4-linked β-Gal*p*A, respectively. In HSQC (**Fig. 2B**) spectrum, signals at δ 4.71/97.49 and δ 5.40/93.52 suggested the correlations of anomeric carbon and hydrogen of 4-linked Gal*p*A **(i, h)**, which indicating that 4-linked Gal*p*A residues are β- and α-configuration, respectively. C-1 signal of T-linked β-Gal*p* **(g)** was overlapped by C-1 signal of 1, 3-linked β-Gal*p* **(f)** and assigned to δ 104.67. Resonance at δ 100.25 could be assigned to C1 of 1, 2, 4-linked α-Rha*p* **(c)** and its correlation with H1 of this residue at δ 5.15 in HSQC spectrum (**Fig. 2B**). Arabian residues were too weak to be directly observed in ^13^C NMR spectrum. However, combined with the cross peaks at higher chemical shift in HSQC spectrum (**Fig. 2B**), resonances at δ 5.28/108.18 and δ 5.17/108.61 could be easily assigned to H1 and C1 of T-linked α-Ara*f* **(a)** and 1, 5-linked α-Ara*f* **(b)**, respectively. From C-2 to C-6 regions, the strong signals at δ 69.11, δ 79.89 and δ 71.83 were allocated to atoms C-2, C-4 and C-5 of residue **d** (1, 4-linked α-Gal*p*A), respectively.

Resonances at δ 69.22, δ 79.44 and δ 71.94 were assigned to atoms C-2, C-4 and C-5 of residue **d’** (1, 4-linked α-Gal*p*A), respectively. Signals at δ 71.81 and δ 83.30 were assigned to atoms C-2 and C-4 of residues **e (**1, 4-linked α-6MeOGal*p*A) and **e’ (**1, 4-linked α-6MeOGal*p*A) in different chemical environments, respectively. Signal at δ 80.27 could belong to C-3 of residue **f (**1, 3-linked β-Gal*p*). In ^1^H NMR (**Fig. 1A**) spectrum of 922211, the anomeric hydrogens at δ 5.15 of **c (**1, 2, 4-linked α-Rha*p***), d (**1, 4-linked α-Gal*p*A) and **d’ (**1, 4-linked α-Gal*p*A) were overlapped heavily. Signals at δ 5.01 could be assigned to H1 of **e (**1, 4-linked α-6MeOGal*p*A) and **e’ (**1, 4-linked α-6MeOGal*p*A). Signals at δ 4.70 originated from H-1 of **g (**T-linked β-Gal*p*) and **f (**1, 3-linked β-Gal*p*), respectively. Resonance at δ 4.71 originated from H-1 of and **i (**4-linked β-Gal*p*A).

Resonance at δ 5.40 could be assigned to H-1 of **h (**4-linked α-Gal*p*A). Signals at δ 5.28 and δ 5.17 could be assigned to H-1 of **a (**T-linked α-Ara*f*) and **b (**1, 5-linked α-Ara*f*), respectively. Comparing with 922211, the resonances belonging to T-, 1, 3-linked Gal and T-, 1, 5-linked Ara almost vanished in HSQC of P2922211I (**Fig. 1C**). Hence, it might be deduced that those vanished glycosyl residues are on side chains.

In HMBC spectrum (**Fig. 3**), strong resonance at δ 5.15/79.89 indicated that the H-1 of **d (**1, 4-linked α-Gal*p*A) was correlated with next C-4 of **d (**1, 4-linked α-Gal*p*A). Meanwhile, the correlation at δ 100.85/4.52 suggested that the C-1 of **d (**1, 4-linked α-Gal*p*A) was correlated with next H-4 of **d (**1, 4-linked α-Gal*p*A). Hence, 1, 4-linked α-Gal*p*A residues are adjacent to each other. The cross peaks at δ 5.15/83.30 and δ 100.85/4.49 indicated that the H-1 of **d/d’** (1, 4-linked α-Gal*p*A/another 1, 4-linked α-Gal*p*A) was correlated with C-4 of **e/e’ (**1, 4-linked α-6MeOGal*p*A/another 1, 4-linked α-6MeOGal*p*A) and the C-1 of **d (**1, 4-linked α-Gal*p*A) was correlated with H-4 of **e’ (**1, 4-linked α-6MeOGal*p*A), respectively. The resonances at δ 4.52/101.67 and δ 79.89/5.01 showed that the H-4 of **d/d’** (1, 4-linked α-Gal*p*A/ another 1, 4-linked α-Gal*p*A) was coupled with C-1 of **e (**1, 4-linked α-6MeOGal*p*A) and the C-4 of **d (**1, 4-linked α-Gal*p*A) was allocated with H-1 of **e/e’ (**1, 4-linked α-6MeOGal*p*A/1, 4-linked α-6MeOGal*p*A), respectively. Cross peaks at δ 5.15/78.97 and δ 100.85/4.20 showed the correlation between H-1 of **d/d’** (1, 4-linked α-Gal*p*A/another 1, 4-linked α-Gal*p*A) and C-2 of **c (**1, 2, 4-linked α-Rha*p*) and association between C-1 of **d (**1, 4-linked α-Gal*p*A) and H-2 of **c (**1, 2, 4-linked α-Rha*p*). Cross peaks at δ 5.15/78.89 and δ 101.08/4.09 suggested the correlation between H-1 of **d/d’** (1, 4-linked α-Gal*p*A/another 1, 4-linked α-Gal*p*A) and C-4 of **c (**1, 2, 4-linked α-Rha*p*) and coupling between C-1 of **d’ (**1, 4-linked α-Gal*p*A) and H-4 of **c (**1, 2, 4-linked α-Rha*p*). Cross peak at δ 5.15/79.89/79.44 suggested the correlation between H-1 of **c (**1, 2, 4-linked α-Rha*p*) was correlated with C-4 of **d/d’** (1, 4-linked α-Gal*p*A/another 1, 4-linked α-Gal*p*A). Cross peak at δ 100.25/4.52 suggested the allocation between C-1 of **c (**1, 2, 4-linked α-Rha*p*) and H-4 of **d/d’** (1, 4-linked α-Gal*p*A/another 1, 4-linked α-Gal*p*A). The resonances at δ 5.15/79.29 and δ 100.25/4.53 showed that the H-1 of **c (**1, 2, 4-linked α-Rha*p*) was correlated with C-4 of **h (**4-linked α-Gal*p*A) and the C-1 of **c (**1, 2, 4-linked α-Rha*p*) was associated with H-4 of **h (**4-linked α-Gal*p*A), respectively. Cross peaks at δ 5.15/78.74 and δ 100.25/4.47 suggested the correlation between H-1 of **c (**1, 2, 4-linked α-Rha*p*) was correlated with C-4 of **i** (4-linked β-Gal*p*A) and the C-1 of **c (**1, 2, 4-linked α-Rha*p*) was coupled with H-4 of **i** (4-linked β-Gal*p*A), respectively. Cross peak at δ 5.17/78.89 suggested the correlation between H-1 of **b (**1, 5-linked α-Ara*f*) was allocated with C-4 of **c (**1, 2, 4-linked α-Rha*p*). Cross peak at δ 5.28/69.84 suggested the correlation between H-1 of **a (**T-linked α-Ara*f*) was correlated with adjacent C-5 of **b (**1, 5-linked α-Ara*f*). Cross peak at δ 4.70/78.89 suggested the correlation between H-1 of **f** (1, 3-linked β-Gal*p*) was correlated with C-4 of **c (**1, 2, 4-linked α-Rha*p*). Cross peaks at δ 4.70/80.27 and δ 104.67/3.73 suggested the correlation between H-1 of **g** (T-linked β-Gal*p*) was correlated with C-3 of **f** (1, 3-linked β-Gal*p*) and the C-1 of **g** (T-linked β-Gal*p*) was correlated with H-3 of **f** (1, 3-linked β-Gal*p*), respectively. Cross peak at δ 5.28/78.89 suggested the correlation between H-1 of **a** (T-linked α-Ara*f*) was correlated with C-4 of **c (**1, 2, 4-linked α-Rha*p*).

**Fig. 3.**
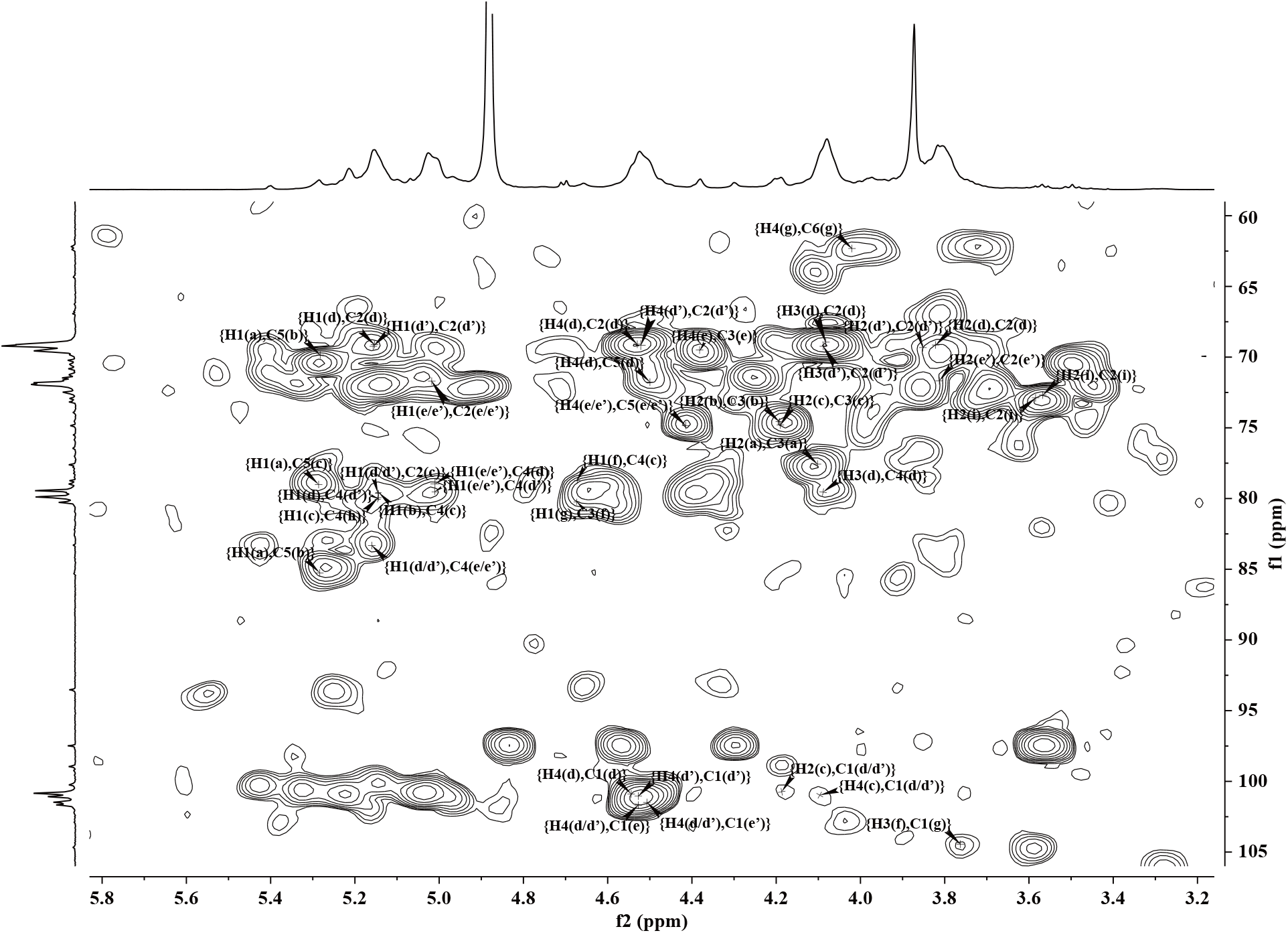
HMBC spectrum of 922211 (**a**. T-linked a-Ara*f*; **b**. 1,5 -linked a-Ara*f*; **c**. 1, 2, 4-linked a-Rha*p*; **d/d’**. 1, 4-linked a-Gal*p*A; **e/e’**. 1, 4-linked α-6MeOGal*p*A; **f**. 1, 3-linked β-Gal*p*; **g**. T-linked β-Gal*p*; **h**. 4-linked a-Gal*p*A; **i**. 4-linked β-Gal*p*A).

Combined with the result mentioned above, the intra-dialysate P2922211I contained 1, 4-linked α-Gal*p*A and 1, 2-linked α-Rha*p* in the molar ratio of 16.71 : 1.00 (47.12 : 2.82 in **table 2**). However, linkage type analysis of the native polysaccharide 922211 showed that it contained 1, 4-linked α-Gal*p*A and 1, 2, 4-linked α-Rha*p* in the molar ratio of 37.91 : 6.40. Based on the huge different proportion of 1, 4-linked α-Gal*p*A and 1, 2-linked α-Rha*p* or 1, 2, 4-linked α-Rha*p* in P2922211I and 922211, it was reasonably deduced that the backbone consisted of two segments: linear homogalacturonan chains which might be partly methyl esterified and RG I-type-like fragment bearing 1, 4-linked α-Gal*p*A, 1, 4-linked α-Gal*p*AOMe and 1, 2, 4-linked α-Rha*p*. There are four branches attached to C-1 or C4 position of Rhamnose glycosyl residues on backbone. Hence, taken together, the proposed repeating unit of 922211 was presented as following:

**Fig S1.**
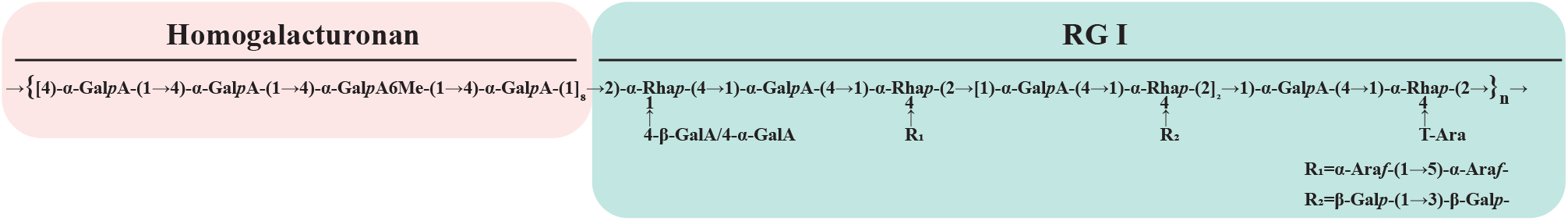
Scheme 1. Schematic structure of 922211.

### 3.5. Crude polysaccharide 922 was a potent inhibitor of SARS-COV-2 3CLpro

As early as in the study of coronavirus it was found that 3CL protein was an attractive target molecule against coronavirus. This is because the functional polypeptides are released from the polyproteins by extensive proteolytic processing during coronavirus complex replication. This is primarily achieved by the 33.1-kD HCoV 229E main proteinase (Mpro), which is frequently also called 3C-like proteinase (3CLpro) (Anand et al., 2003). Since the outbreak of COVID-19 in 2019, with the in-depth study of COVID-19, scientists have also found that inhibiting the activity of 3CL protein and disturbing the interaction between SARS-CoV-2-S1 and ACE2 are two feasible strategies for the development the drugs and vaccines in COVID-19. Hence, some compounds like flavonols, Genkwanine, and Luteolin-glucoside have high affinity with ACE2 and 3CLpro (Mouffouk et al., 2021; Nouadi et al., 2021). Indeed, some of their SARS-CoV-2 antiviral effects had also been proved (Mouffouk et al., 2021). Moreover, it was reported that macromolecular carbohydrate such as heparin might block SARS-CoV-2 binding and infection because negatively charged sulfate and carboxyl groups on heparin could stabilized the association with several positively charged amino acid residues of spike protein (Batiha et al., 2020). Our recent study one crude polysaccharide 375 in which contain alginate might potently inhibit SARS-CoV-2 virus replication (Zhang et al., 2021). As we know alginate has β-_D_-mannuronate (M) and 1, 4-linked α-_L_-guluronate. This also suggests that polysaccharide contains uronic acid group may benefit the effect against the virus. However, the most common type of pectic polysaccharides, with a large carboxyl group, has not been reported on COVID-19 yet. Therefore, in this study, a polysaccharide containing a large amount of uronic acid was chosen to screen its activity of inhibiting 3CL protein. Based on the basic characteristics that 2019-nCoV 3CLpro protein is a proteolytic enzyme, a screening system for fluorescence detection of 2019-nCoV 3CLpro protein activity was established. 2019-nCoV 3CLpro protein can specifically shear the substrate with GLN (q) at P1 position. Fluorescent polypeptide can be used as the substrate for its activity detection, and the activity of 3CLpro protein hydrolase can be reflected by detecting the generation of fluorescent signal. Then the competitive binding test targeting 3CLpro was examined. The results showed that the crude 922 and the homogeneous polysaccharide 922211, derived from 922, might potently inhibit SARS-CoV-2 3CLpro activity (**Fig. 4A and B**). Further, the fluorescence resonance energy transfer (FRET) based cleavage assay was employed to determine the median inhibitory concentration (IC_50_) values. The results revealed good inhibitory potency of 922 and 922211, with IC_50_ values of 4.73 ± 1.05 µM and 0.18 ± 0.01 µM (**Fig. 4A and B**), respectively. The above results imply that polysaccharide, as the main component of 922 and 922211, might have a blocking effect on SARS-CoV-2 replication and infection. This inspires us to furtherly explore polysaccharides against SARS-Cov-2 and their underlying mechanism.

**Fig. 4.**
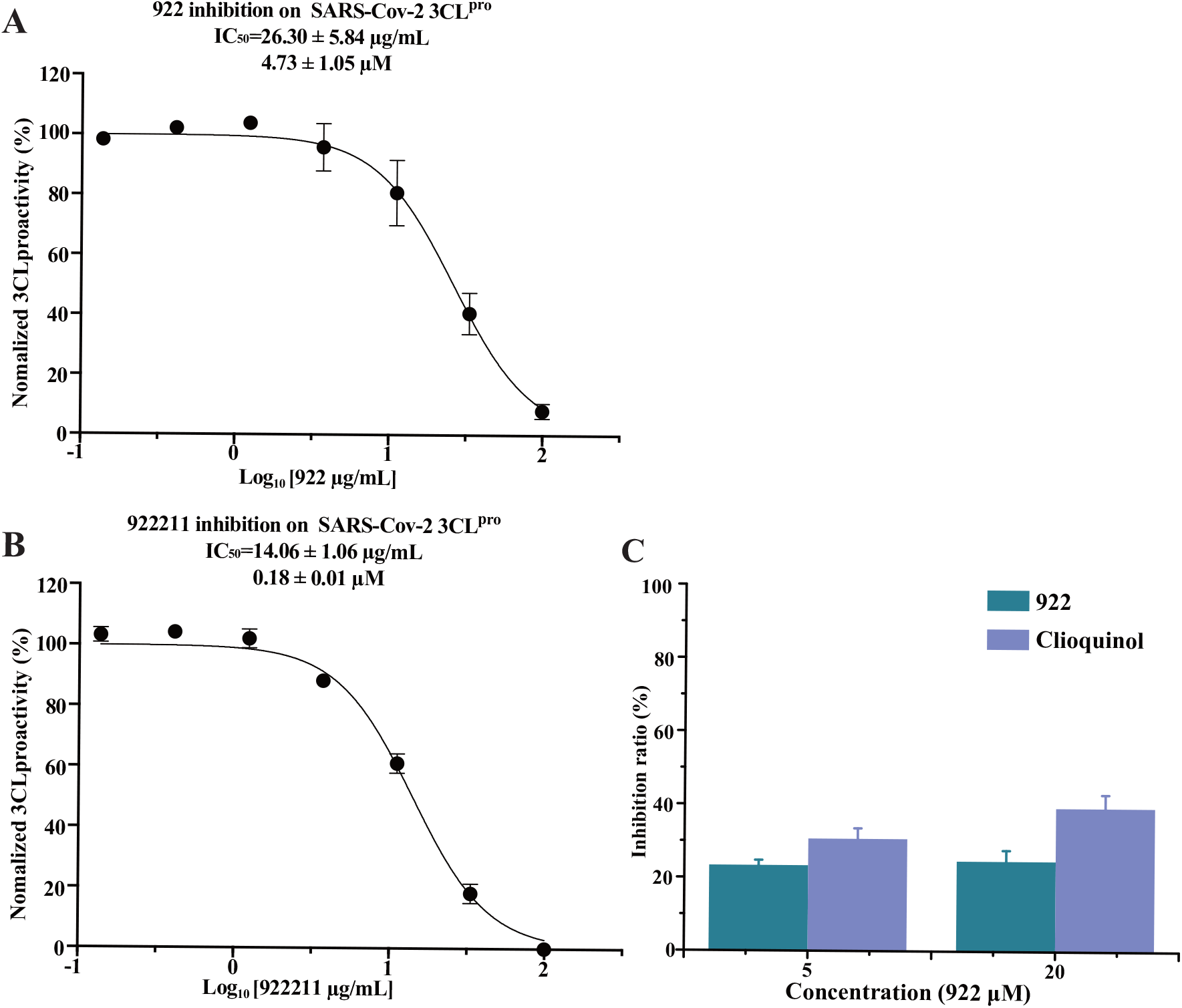
922 and 922211 inhibit the activity of SARS-Cov-2 3CLpro (A, B); Competitive intervention of polysaccharide 922 on S1 protein and ACE2 (C). The protease activity of SARS-CoV-2 3CLpro was measured in the presence of increasing concentrations of the 922 and 922211, SARS-CoV-2 3CLpro preincubated for 20 min with each concentration of 922 and 922211. The protease activity was measured by the FRET-based protease assay. Dose–response curves for IC_50_ values were determined by nonlinear regression. All data are shown as mean ± SD. n = 3 biological replicates (A, B). Competitive intervention of crude polysaccharide 922 and Clioquinol with concentrations of 5 μM and 20 μM on S1 protein and ACE2 interaction by ELISA experiment (C). Final concentrations of ACE2 and biotinylated -S1 protein were 2 μg/mL and 500 ng/mL, respectively.

### 3.6. 922 may disturb the interaction between SARS-CoV-2-S1 and ACE2

The spike protein of SARS-CoV-2 shows more than 90% amino acid similarity to that of pangolin and bat CoVs which also bind to human angiotensin-convert enzyme 2 (ACE2) for the virus infection. Thus, S protein is very vital for viral invasion. Interestingly, unoccupied glycosylation sites on SARS-CoV-2 S and the glycan binding site of N-terminal domain of S1 protein imply S protein may bind with carbohydrate (Kirchdoerfer et al., 2016; Watanabe et al., 2020). To further screen the bioactivity of 922 and 922211 against SARS-Cov-2, ELISA was used to examine whether 922 and 922211 might disturb the binding between S1 protein *in vitro*. The results showed that polysaccharide 922 could impede weakly the binding of S1 protein with ACE2 (**Fig. 4C**). These results also suggest that anti-SARS-CoV-2 effects of 922 and 922211 are more likely to inhibit 3CL protein activity than disturbing the interaction between SARS-CoV-2-S1 and ACE2.

### 3.7. 922 exhibit anti-viral effect on SARS-CoV-2

The above results implied that 922 might have potential inhibitory effect on SARS-Cov-2. Hence, the inhibition effect of native polysaccharide 922 against the virus was examined. Surprisingly, polysaccharide 922 nearly completely blocked SARS-Cov-2 replication *in vitro*, exhibiting very good anti-SARS-CoV-2 activity in Vero E6 cells. The inhibition rate was 99.9% (**Fig. 5A**). The EC_50_ value was 0.90 µM (or 5.01 µg/mL) (**Fig. 5B**). The above results suggested that 922 is the good candidate for anti-SARS-Cov-2 new drug development. Following, we will further explore the antiviral mechanism for 922 and 922211.

**Fig. 5.**
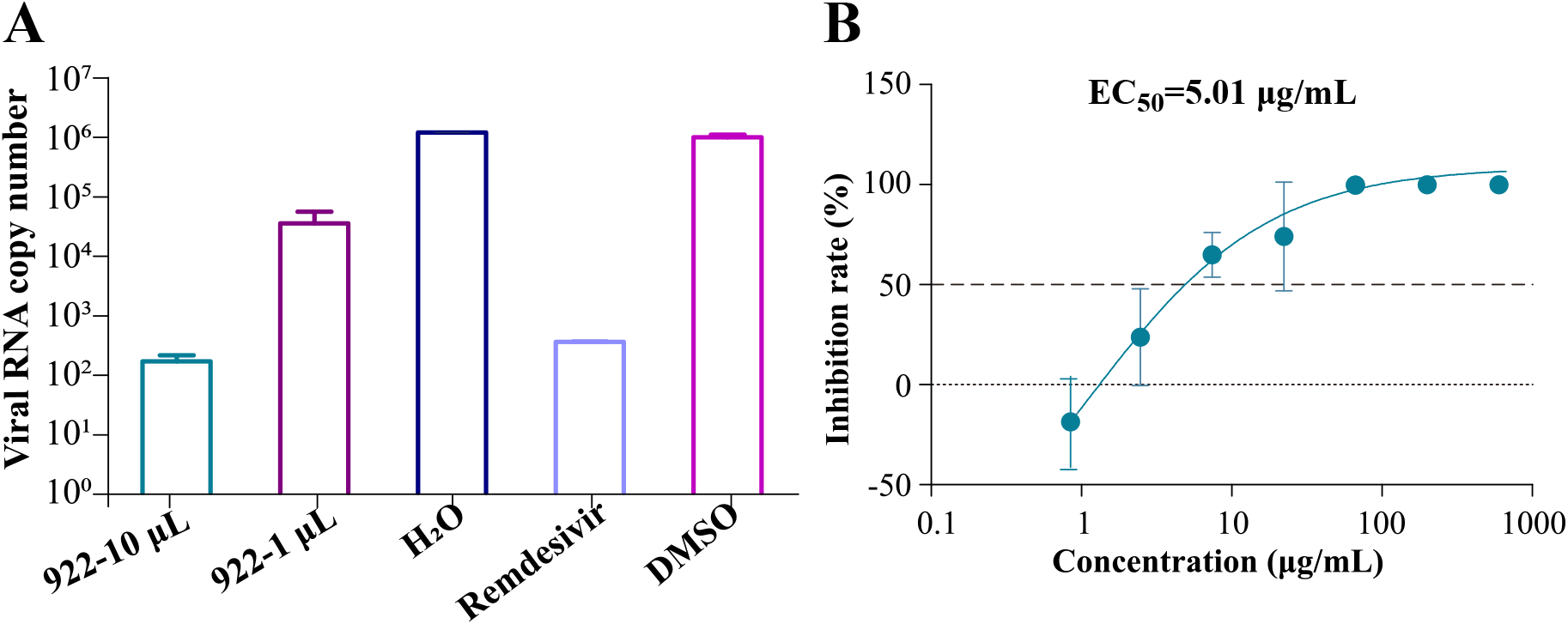
*in vitro* inhibition of polysaccharides against SARS-CoV-2. (A) viral RNA copy number was detected by qPCR after the treatment of solvent (H_2_O) control, Remdesivir positive control (10 μM), crude polysaccharide 922 in 20 μg/mL and 200 μg/mL, respectively. (B) EC_50_ of crude polysaccharide 922 against SARS-CoV-2.

## Discussion

In this study, we uncovered for the first time that polysaccharide and a novel pectin extracted from traditional Chinese medicine *Syzygium aromaticum* flower inhibited strongly the replication of SARS-CoV-2, and the inhibition rate was 99.9%. Further study showed that 922 and 922211 might competitively bind to SARS-CoV-2 key enzyme 3CLpro. The experimental results evidenced our hypothesis that acidic polysaccharides such as pectin might have anti-SARS-CoV-2 effect. In the current anti-virus investigations of polysaccharides, heparin (Gupta et al., 2021; Tandon et al., 2021) or polysaccharides extracted from seaweed (Rosales-Mendoza et al., 2020), alginate (Zhang et al., 2021), with large amounts of sulfate ions and chitosan derivatives (Modak et al., 2021) have been studied. Moreover, the chitosan derivatives are also sulfated chitosan, which has significant or potent effect on anti-SARS-CoV-2. Unlike other sulfated polysaccharides, which antiviral activity is mainly arisen from competitive inhibition of S1 and ACE2 binding, both 922 and the pectin-like glycan 922211 might impede 3CLpro activity, but only tenderly disturb the binding of S1 and ACE2.

To further understand whether there some other components in the 922 may interfere 3CLpro, PLpro, or RdRp enzyme activity. We firstly pooled the fragment from 922 crude polysaccharide eluted by distilled water, 0.05 M NaCl, 0.1 M NaCl, 0.2 M NaCl, 0.4 M NaCl, stepwise to achieve 9220S, 9220P, 92205S, 92205P, 9221S, 9221P, 9222S, 9222P, 9224S, and 9224P, respectively (**Fig. 6F**). Then all of those fragments were employed to test their effects on 3CLpro, PLpro and RdRp enzymes activities. Interestingly, 9220P, 9221P, 9222P, 9224P, 9221S might significantly impede 3CLpro activity (**Fig. 6A-E**). The IC_50_ of 9220P, 9221P, 9222P, 9224P, 9221S on 3CLpro was roughly 76.57 μg/mL, 42.12 μg/mL, 17.17 μg/mL, 14.36 μg/mL, 22.99 μg/mL (**Fig. 6A-E**), respectively. Surprisingly, 92205P, 9222P and 9224P could also potently inhibit RdRp enzyme activity, while 9222P and 9224P might nearly completely block this enzyme activity (**Table S1**). However, almost all of these components have no significant effect on the PLpro enzyme (**Table S1**). This imply that polysaccharide 922 inhibit SARS-CoV-2 virus replication at least through disturbing both 3CLpro and RdRp enzymes, but has nothing to do with PLpro enzyme.

**Fig. 6.**
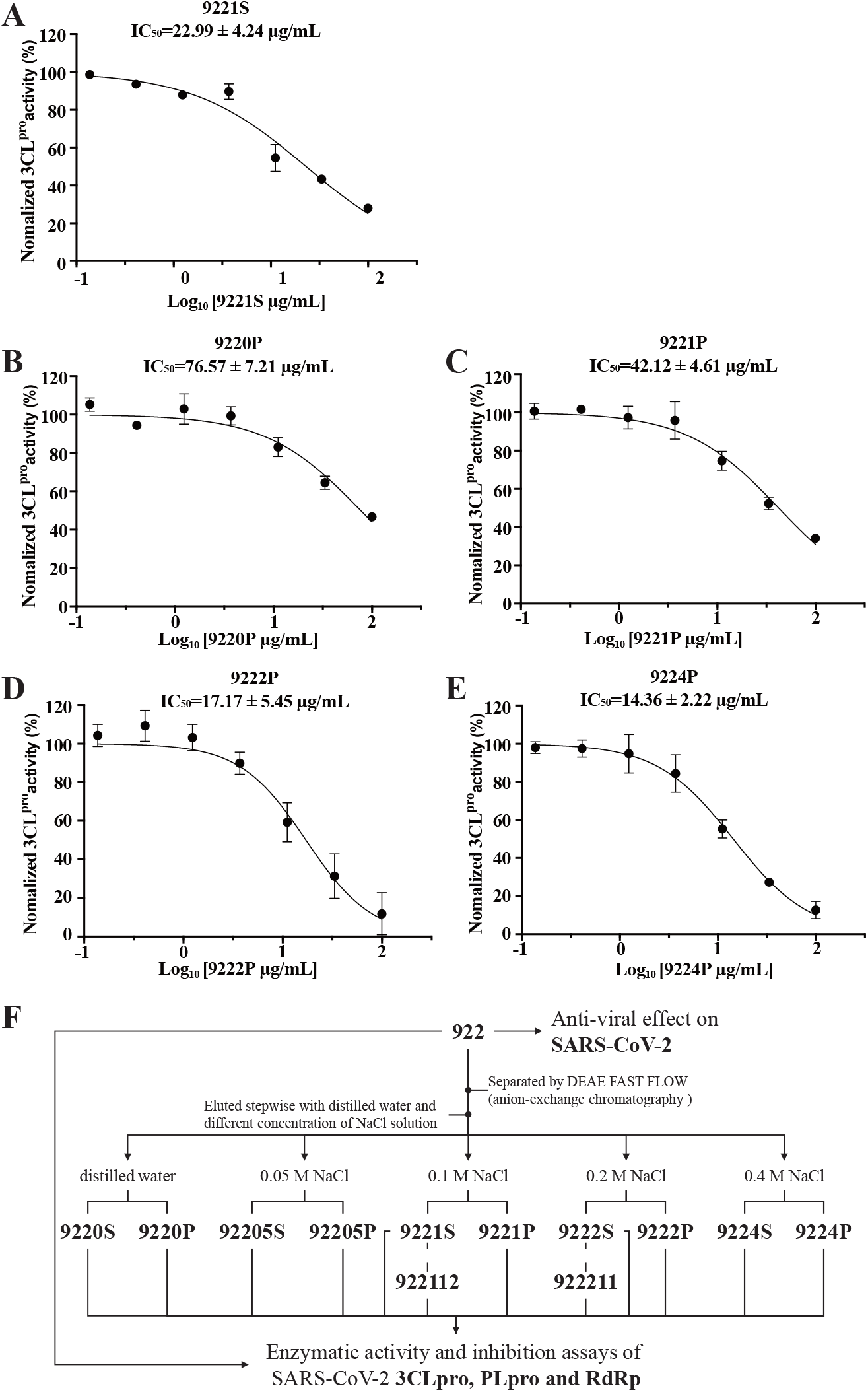
9221S, 9220P, 9221P, 9222P and 9224P inhibit the activity of SARS-Cov-2 3CLpro (A-E). Schematic diagram of 922 separation, purification. 922 and fractions arising from it determinate enzymatic activity and inhibition assays of SARS-CoV-2 3CLpro, PLpro and RdRp (F). The protease activity of SARS-CoV-2 3CLpro was measured in the presence of increasing concentrations of the 9221S, 9220P, 9221P, 9222P and 9224P, SARS-CoV-2 3CLpro preincubated for 20 min with each concentration of above five samples, the protease activity was measured by the FRET-based protease assay. Dose–response curves for IC_50_ values were determined by nonlinear regression. All data are shown as mean ± SD. n = 3 biological replicates. (9220P: the fraction from 922 mainly containing protein separated by DEAE Sepharose Fast Flow (anion-exchange chromatography, eluted by distill water); 9220S: the fraction from 922 mainly containing sugar separated by DEAE Sepharose Fast Flow, eluted by distill water; 92205P: the fraction from 922 mainly containing protein separated by DEAE Sepharose Fast Flow, eluted by 0.05 M NaCl; 92205S: the fraction from 922 mainly containing sugar separated by DEAE Sepharose Fast Flow, eluted by 0.05 M NaCl; 9221P: the fraction from 922 mainly containing protein separated by DEAE Sepharose Fast Flow, eluted by 0.1 M NaCl; 9221S: the fraction from 922 mainly containing sugar separated by DEAE Sepharose Fast Flow, eluted by 0.1 M NaCl; 9222P: the fraction from 922 mainly containing protein separated by DEAE Sepharose Fast Flow, eluted by 0.2 M NaCl; 9222S: the fraction from 922 mainly containing sugar separated by DEAE Sepharose Fast Flow, eluted by 0.2 M NaCl; 9224P: the fraction from 922 mainly containing protein separated by DEAE Sepharose Fast Flow, eluted by 0.4 M NaCl; 9224S: the fraction from 922 mainly containing sugar separated by DEAE Sepharose Fast Flow, eluted by 0.4 M NaCl; 9222111: the homogenous polysaccharide from 9222S purified by Sephacryl S-100 HR (gel permeation chromatography); 922111: the polysaccharide from 9221S purified by Sephacryl S-100 HR).

Although the detail mechanism underlying the action of 922 and 922211 need to be further explored, our study provides evidences for the first time for pectin or pectin-like polysaccharide might be promising candidate for the anti-SARS-CoV-2 virus new drug development.

## Supporting information

Supplementary datas

## Acknowledgment

This work was supported by Shanghai Municipal Science and Technology Major Project, Key New Drug Creation and Manufacturing Program (Grant number 2019ZX09735001), National Key R&D Program, Ministry of Science and Technology of China (Grant number 2019YFC1711-000), COVID-19 Emergency Research Project founded by Zhejiang University (2020XGZX080). We are particularly grateful to Tao Du and Lun Wang from Zhengdian Biosafety Level 3 Laboratory and the running team of the laboratory for their work

