## Supplementary datas for "Novel pectin from crude polysaccharide of *Syzygium aromaticum* against SARS-CoV-2 activities by targeting 3CLpro"

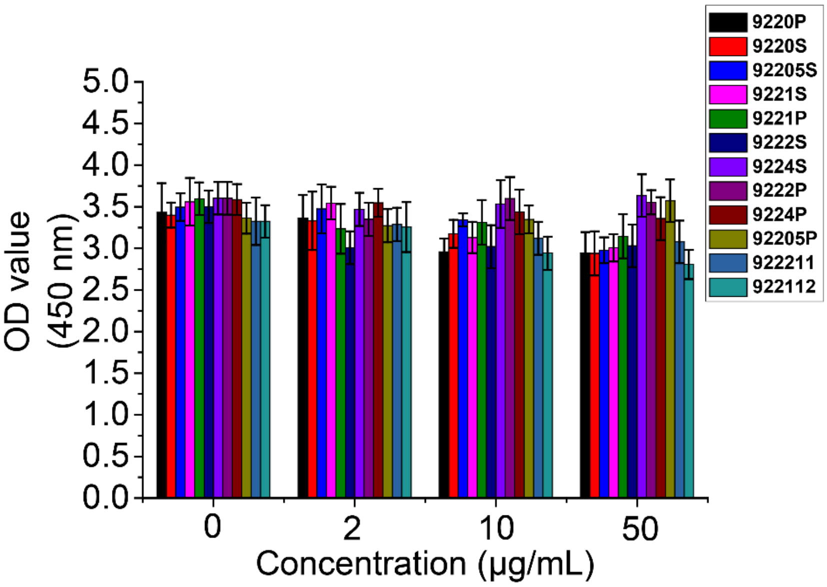


**Figure S1.** Inhibitory activity profiles of 12 samples (0, 2, 10, 50 μg/mL) separated and/or purified from 922 against ACE2 and S1 protein binding by ELISA. OD value means the affinity between ACE2 and S1 protein. The final used concentrations of ACE2 and biotinylated -S1 protein were 2 μg/mL and 500 ng/mL, respectively. (9220P: the fraction from 922 mainly containing protein separated by DEAE Sepharose Fast Flow (anion-exchange chromatography, eluted by distill water); 9220S: the fraction from 922 mainly containing sugar separated by DEAE Sepharose Fast Flow, eluted by distill water; 92205P: the fraction from 922 mainly containing protein separated by DEAE Sepharose Fast Flow, eluted by 0.05 M NaCl; 92205S: the fraction from 922 mainly containing sugar separated by DEAE Sepharose Fast Flow, eluted by 0.05 M NaCl; 9221P: the fraction from 922 mainly containing protein separated by DEAE Sepharose Fast Flow, eluted by 0.1 M NaCl; 9221S: the fraction from 922 mainly containing sugar separated by DEAE Sepharose Fast Flow, eluted by 0.1 M NaCl; 9222P: the fraction from 922 mainly containing protein separated by DEAE Sepharose Fast Flow, eluted by 0.2 M NaCl; 9222S: the fraction from 922 mainly containing sugar separated by DEAE Sepharose Fast Flow, eluted by 0.2 M NaCl; 9224P: the fraction from 922 mainly containing protein separated by DEAE Sepharose Fast Flow, eluted by 0.4 M NaCl; 9224S: the fraction from 922 mainly containing sugar separated by DEAE Sepharose Fast Flow, eluted by 0.4 M NaCl; 9222111: the homogenous polysaccharide from 9222S purified by Sephacryl S-100 HR (gel permeation chromatography); 922111: the polysaccharide from 9221S purified by Sephacryl S-100 HR (gel permeation chromatography).

**Table S1.** Enzymatic activity and inhibition assays of SARS-CoV-2 3CLpro, PLpro and RdRp of 922 and fractions arising from it.

| | Seq | Compound Name | Inhibition Rate% (100 μg/mL) | $IC_{50}$ ( μg/mL) |
|---|---|---|---|---|
| The enzyme activity and inhibition assays of SARS-CoV-2 3CLpro | 1 | 922 | 91.8±2.5 | 26.30±5.84 |
| | 2 | 9220S | 15.8±12.2 | —— |
| | 3 | 92205S | 14.9±8.3 | —— |
| | 4 | 9221S | 72.1±1.7 | 22.99±4.24 |
| | 5 | 9222S | 0.0±0.0 | —— |
| | 6 | 9224S | -3.1±15.5 | —— |
| | 7 | 9220P | 53.2±6.9 | 76.57±7.21 |
| | 8 | 92205P | 35.3±4.3 | —— |
| | 9 | 9221P | 65.8±1.4 | 42.12±4.61 |
| | 10 | 9222P | 88.2±10.9 | 17.17±5.45 |
| | 11 | 9224P | 87.2±4.5 | 14.36±2.22 |
| | 12 | 922112 | 113.5±1.5 | 7.72±0.65 |
| | 13 | 922211 | -8.4±11.5 | —— |
| The enzyme activity and inhibition assays of SARS-CoV-2 PLpro | 1 | 922 | 91.4±1.3 | 19.72±2.20 |
| | 2 | 9220S | 4.1±2.2 | —— |
| | 3 | 92205S | 4.4±3.7 | —— |
| | 4 | 9221S | 2.8±1.8 | —— |
| | 5 | 9222S | 0.0±0.0 | —— |
| | 6 | 9224S | 0.0±0.0 | —— |
| | 7 | 9220P | 5.1±1.7 | —— |
| | 8 | 92205P | 0.0±0.0 | —— |
| | 9 | 9221P | 0.0±0.0 | —— |
| | 10 | 9222P | 0.0±0.0 | —— |
| | 11 | 9224P | 0.0±0.0 | —— |
| | 12 | 922112 | 1.5±0.5 | —— |
| | 13 | 922211 | 0.0±0.0 | —— |
| The enzyme activity and inhibition assays of SARS-CoV-2 RdRp | 1 | 922 | —— | —— |
| | 2 | 9220S | 20.8±0.4 | —— |
| | 3 | 92205S | 7.3±5.6 | —— |
| | 4 | 9221S | 20.8±0.8 | —— |
| | 5 | 9222S | 5.6±2.3 | —— |
| | 6 | 9224S | 0.0±0.0 | —— |
| | 7 | 9220P | 0.0±0.0 | —— |
| | 8 | 92205P | 35.3±7.3 | —— |
| | 9 | 9221P | 3.7±3.6 | —— |
| | 10 | 9222P | 92.8±2.2 | 50.29±9.48 |
| | 11 | 9224P | 104.3±0.9 | 62.99±4.38 |
| | 12 | 922112 | 0.0±0.0 | —— |
| | 13 | 922211 | 0.0±0.0 | —— |
| | | | | $IC_{50}$ ( μM) |
| | Control group | Remdesivir triphosphate | | 1.80±0.25 |
| | | Suramin | | 0.54±0.17 |

9220P: the fraction from 922 mainly containing protein separated by DEAE Sepharose Fast Flow (anion-exchange chromatography, eluted by distill water); 9220S: the fraction from 922 mainly containing sugar separated by DEAE Sepharose Fast Flow, eluted by distill water; 92205P: the fraction from 922 mainly containing protein separated by DEAE Sepharose Fast Flow, eluted by 0.05 M NaCl; 92205S: the fraction from 922 mainly containing sugar separated by DEAE Sepharose Fast Flow, eluted by 0.05 M NaCl; 9221P: the fraction from 922 mainly containing protein separated by DEAE Sepharose Fast Flow, eluted by 0.1 M NaCl; 9221S: the fraction from 922 mainly containing sugar separated by DEAE Sepharose Fast Flow, eluted by 0.1 M NaCl; 9222P: the fraction from 922 mainly containing protein separated by DEAE Sepharose Fast Flow, eluted by 0.2 M NaCl; 9222S: the fraction from 922 mainly containing sugar separated by DEAE Sepharose Fast Flow, eluted by 0.2 M NaCl; 9224P: the fraction from 922 mainly containing protein separated by DEAE Sepharose Fast Flow, eluted by 0.4 M NaCl; 9224S: the fraction from 922 mainly containing sugar separated by DEAE Sepharose Fast Flow, eluted by 0.4 M NaCl; 9222111: the homogenous polysaccharide from 9222S purified by Sephacryl S-100 HR (gel permeation chromatography); 922111: the polysaccharide from 9221S purified by Sephacryl S-100 HR.
